# coelsch: Platform-agnostic single-cell analysis of meiotic recombination

**DOI:** 10.64898/2026.02.02.701279

**Authors:** Matthew T. Parker, Samija Amar, Jonas Freudigmann, Birgit Walkemeier, Xiao Dong, Victor Solier, Magdalena Marek, Bruno Huettel, Raphael Mercier, Korbinian Schneeberger

## Abstract

**Background:** Meiotic recombination creates genetic diversity through reciprocal exchange of haplotypes between homologous chromosomes. Scalable and robust methods for mapping recombination breakpoints are essential for understanding meiosis and for genetic mapping. Single cell sequencing of gametes offers a direct approach to recombination mapping, yet the effect of technical differences between single-cell sequencing methods for crossover detection remains unclear.

**Results:** We benchmark single cell methods for droplet-based chromatin accessibility and RNA sequencing and plate-based whole-genome amplification for mapping meiotic recombination in *Arabidopsis thaliana*. For this purpose we introduce two novel open-source tools **coelsch_mapping_pipeline** and **coelsch** for haplotype-aware alignment and per-cell crossover detection, using them to recover known recombination frequencies and quantify the effects of coverage sparsity. We subsequently apply our approach to a panel of 40 recombinant F₁ hybrids derived from crosses of 22 diverse natural accessions, successfully recovering genetic maps for 34 F_1_s in a single dataset. This analysis reveals substantial variation in recombination rate and identifies a ∼10 Mb pericentric inversion in the accession Zin-9, the largest natural inversion reported in *A. thaliana* to date.

**Conclusions:** These results demonstrate the applicability and scalability of single-cell gamete sequencing for high-throughput mapping of meiotic recombination, and highlight the strengths and limitations of different single-cell modalities. The accompanying open-source tools provide a framework for haplotyping and crossover detection analysis using sparse single-cell sequencing data. Our methodology enables parallel analysis of large numbers of hybrids in a single dataset, removing a major technical barrier to large-scale studies of natural variation in recombination rate.

## Background

Meiotic recombination generates genetic diversity by shuffling parental alleles through the exchange of chromosome arms (i.e., crossovers) (1). The frequency and distribution of crossovers along the genome varies widely across species, individuals, and genomic regions (2). This variation is shaped by both *cis*-acting factors such as chromatin state and sequence divergence, and *trans*-acting variation in the meiotic machinery itself (3–6). Understanding how these factors interact to shape recombination landscapes is a key question in molecular biology (1), and has great significance for evolutionary processes such as mutation rate, population dynamics, hybridisation, and speciation (2). In order to study meiosis, however, accurate genome-wide maps of crossover locations are required (7–9).

Traditional approaches for mapping crossovers rely on sequencing large populations of recombinant progeny (9). While effective, these methods are labour-intensive, and most work has been limited to a handful of genetic backgrounds (9–11). Where applicable, single-cell sequencing of gametes provides a credible high-throughput alternative. Because each gamete is a direct product of a single meiosis, haplotyping individual pollen nuclei allows reconstruction of recombinant chromosomes without progeny generation (12). However, different single-cell modalities sample distinct molecular fractions - RNA, accessible chromatin, or genomic DNA - with varying coverage, noise characteristics, and doublet rates (where two or more cells are captured as a single barcode) (13). The consequences of these technical differences for haplotype inference and crossover detection remain largely unquantified.

Several approaches for haplotyping of recombinant pollen have previously been introduced, based either on single-molecule long-read sequencing (7,14) or sequencing of individual gamete genomes (12,15–18), and demonstrating that crossover landscapes can be recovered from sparse sequencing data. Building on this foundation, we sought to generalise the approach across commonly used single-cell sequencing platforms and modalities. To this end, we have developed two open-source software tools: **coelsch_mapping_pipeline**, which performs haplotype-aware alignment of single cell sequencing reads across parental assemblies; and **coelsch**, which implements per-cell data cleaning and crossover detection via a Hidden Markov Model.

Here, we apply these tools to single-nucleus datasets from *Arabidopsis thaliana* pollen generated using droplet-based RNA and chromatin accessibility assays, as well as plate-based whole-genome amplification. We test the ability of each modality to recover known recombination frequencies, including in mutants with altered crossover rates, and use simulations to assess the impact of coverage and signal sparsity on detection sensitivity. Finally, we analyse 34 recombinant F₁ hybrids generated by crossing 19 different natural accessions, showcasing the power of analysing variation in recombination rate and distribution across a diverse panel. This analysis identifies both trans-acting effects and local structural variants shaping the Arabidopsis recombination landscape, including a previously unrecognised 10 Mb pericentric inversion in the Zin-9 accession. Together, these results demonstrate how high-throughput single-cell sequencing can be applied to study meiotic recombination at scale.

## Results

### Comparison of single-cell sequencing modalities for recombination analysis

To evaluate how different single-cell sequencing methods perform in the mapping of meiotic recombination, we generated datasets from Arabidopsis Col-0 × L*er* F₁ pollen using four modalities: 10x Chromium single cell accessible chromatin sequencing (10x scATAC), BD Rhapsody and 10x Chromium 3′ single cell RNA-seq (BD scRNA, 10x scRNA), and Takara iCELL8 single cell whole genome amplification (Takara scWGA). Each platform samples a different molecular fraction and exhibits distinct trade-offs between number of cells/nuclei and per-cell coverage (Figure 1A).

**Figure 1.**
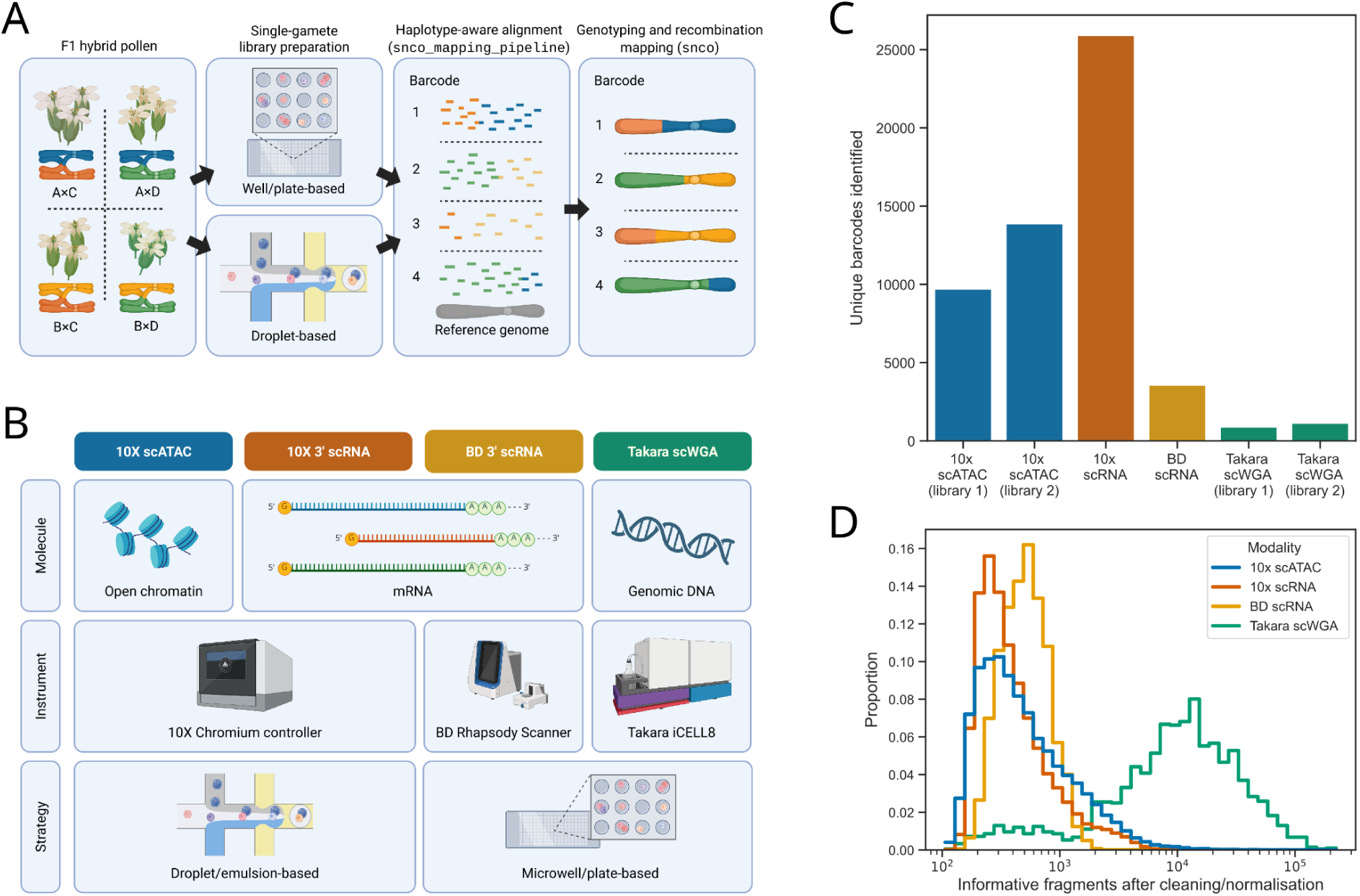
Overview of single-cell sequencing modalities and data characteristics for recombination analysis. **(a)** Experimental workflow for crossover detection in recombinant *Arabidopsis thaliana* pollen. F₁ hybrids were generated and pollen nuclei isolated for droplet-or plate-based single-cell sequencing. Reads were aligned to parental genomes to perform parental genotyping and reconstruct recombinant haplotypes. **(b)** Comparison of the four sequencing modalities used in this study, showing target molecule, instrument, and capture strategy: 10x scATAC (open chromatin, 10x Chromium), 10x scRNA (mRNA, 10x Chromium), BD scRNA (mRNA, BD Rhapsody), and Takara scWGA (genomic DNA, iCELL8). **(c)** Number of unique barcodes detected per library for each modality, after sequencing and initial quality control. **(d)** Distribution of informative fragments per barcode, illustrating differences in per-nucleus coverage among modalities.

To process these diverse datasets, we developed two complementary tools. The first, **coelsch_mapping_pipeline**, performs variant-aware alignment across multiple parental haplotypes using STAR consensus (19,20) and generates haplotype-tagged BAMs for downstream analysis, simultaneously processing single cell barcodes and unique molecular identifiers. The second, **coelsch**, performs complete single-cell crossover analysis from aligned reads or variant count matrices (21), including genotype assignment, background correction, doublet filtering, and HMM-based haplotyping. Together they form a unified framework for comparing recombination landscapes across sequencing modalities (Figure 1B).

After basic coverage filtering, the droplet-based 10x and BD libraries yielded thousands to tens of thousands of nuclei, with hundreds of fragments per barcode that are informative in distinguishing the two haplotypes, whereas the plate-based Takara scWGA contained only around one thousand barcodes per library. However, sequencing DNA instead of RNA or open chromatin achieved an order-of-magnitude higher per-barcode coverage in the Takara scWGA libraries (Figure 1C-D; Table 1). These contrasts demonstrate the expected balance between breadth and depth: droplet methods (scATAC, scRNA) enable population-level recombination profiling, while Takara scWGA enables higher resolution crossover detection in fewer nuclei.

**Table 1.**
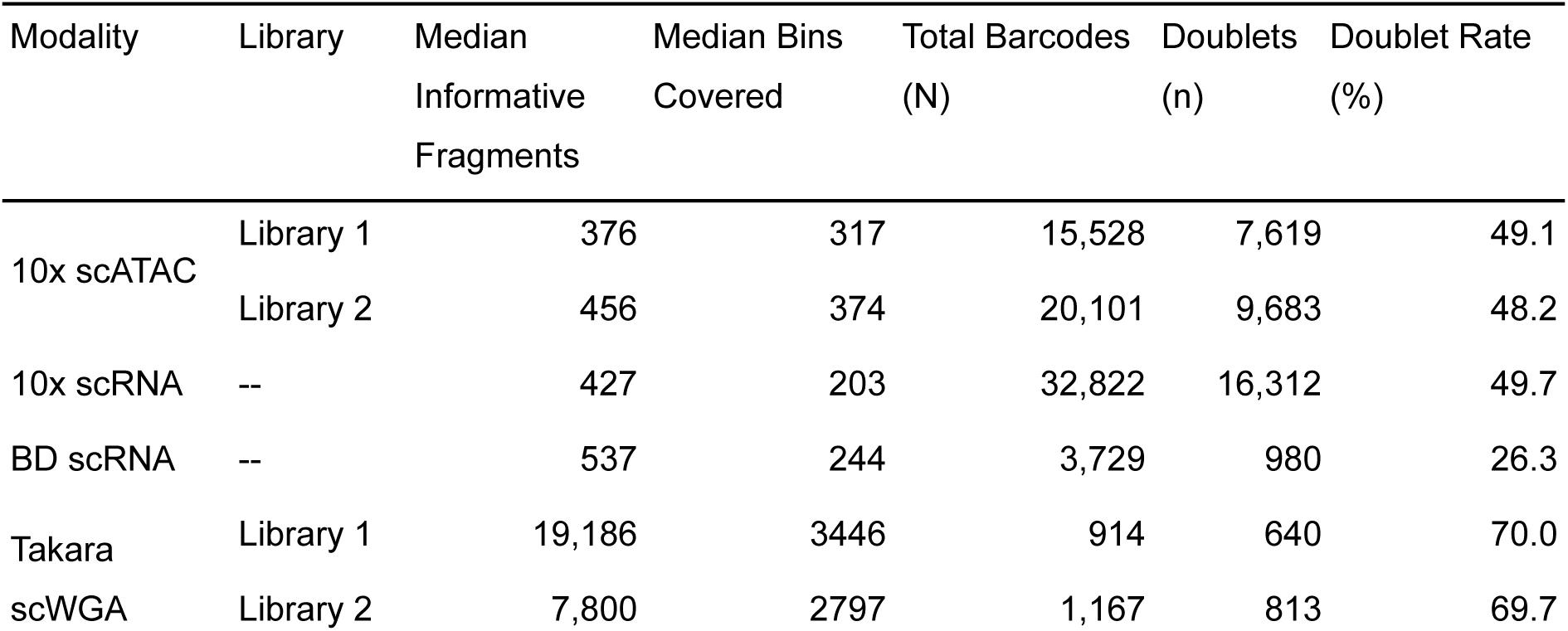
Summary of doublet detection across single-cell modalities. Total barcodes passing quality control (retaining >300 informative marker fragments). Median informative fragments are calculated before data cleaning/normalisation (see Methods). Median bins covered describes the median per-barcode number of 25 kb genomic bins containing at least one informative fragment (out of 5,335 total bins). Doublet classifications are based on a doublet probability threshold of >0.5.

Not only the number of informative fragments, but also their genomic distribution, affects crossover calling. Because 10x and BD scRNA-seq sample mRNA, reads are confined to expressed genes, leaving centromeres and other gene-poor regions sparsely covered. Since the expression of genes ranges over several orders of magnitude, a small number of highly expressed genes can also contribute a disproportionate number of informative fragments. As a result, genomic coverage of scRNA data is less even than scATAC or scWGA. This is evident when counting 25 kb bins containing at least one informative fragment: despite similar median numbers of informative fragments per barcode, 10x scATAC barcodes cover on average 1.7× as many bins as 10x scRNA barcodes (Table 1).

### Detection and filtering of barcode doublets

When two or more cells or nuclei enter the same droplet or well, they are captured under a single molecular barcode, producing a doublet artefact. Because every pollen nucleus carries a unique combination of recombinant haplotypes, doublets will appear heterozygous or noisy. The presence of reads supporting multiple haplotypes within the same genomic region can therefore be used to predict and remove doublets. Coelsch implements a data-driven doublet classifier inspired by scRNA-seq workflows: synthetic doublets are simulated by combining informative reads from multiple barcodes, and mixture modelling analysis of summary metrics identifies real barcodes with doublet-like profiles. Using this approach, we identified high rates of barcode doublets in 10x scATAC (48.6%), 10x scRNA (49.7%) and Takara scWGA (69.8%) datasets (Table 1).

The elevated doublet rate in 10x scATAC data likely stems from the small input volumes required by the Chromium controller, which necessitates centrifugation that can cause clumping and bursting of nuclei. The input volume for 10x scRNA is larger, meaning centrifugation can be avoided, and the high doublet rate observed in the 10x scRNA dataset is inconsistent with our prior experience using this modality (12) and may be an outlier. Unexpectedly, the Takara iCELL8 platform also showed a very high doublet rate. Although the system includes microscopy-based automated well selection which is intended to prevent the dispensing and sequencing of wells containing doublets, 80% of wells classified as *Good* were identified by coelsch as doublets, indicating that the imaging software frequently mis-labels small pollen nuclei (Supplementary Figure 1). Future versions of the platform may benefit from revised image-classification thresholds or omitting the automated selection step altogether.

### Validation of recombination detection across modalities and mutants

Among the Takara scWGA set, 224 high-quality barcodes were recovered from wild-type Col-0 × L*er* pollen and 404 from recombination mutants (*zyp1*, *figl1*, *recq4ab*, *recq4ab zyp1*, *recq4ab figl1*). Crossover detection with coelsch yielded median counts of five crossovers per nucleus for wild-type 10x scATAC and Takara scWGA datasets respectively (Figure 2A), consistent with expected male meiotic rates in Arabidopsis (10). Recovered crossover rates in both 10x scRNA and BD scRNA datasets were lower, likely reflecting fewer and less evenly distributed informative fragments in these datasets (Figure 2A). Mutant datasets (Takara scWGA) were consistent with previously reported effects: *figl1* and *zyp1* showed modest alterations, whereas *recq4ab* displayed a strong elevation, further increased in the double mutants *recq4ab figl1* and *recq4ab zyp1* (Figure 2B). Genomic distributions of crossovers in mutants also approximated previously identified patterns (Figure 2C) (11). These results confirm that single-cell haplotyping can correctly measure quantitative differences in recombination rate across genotypes.

**Figure 2.**
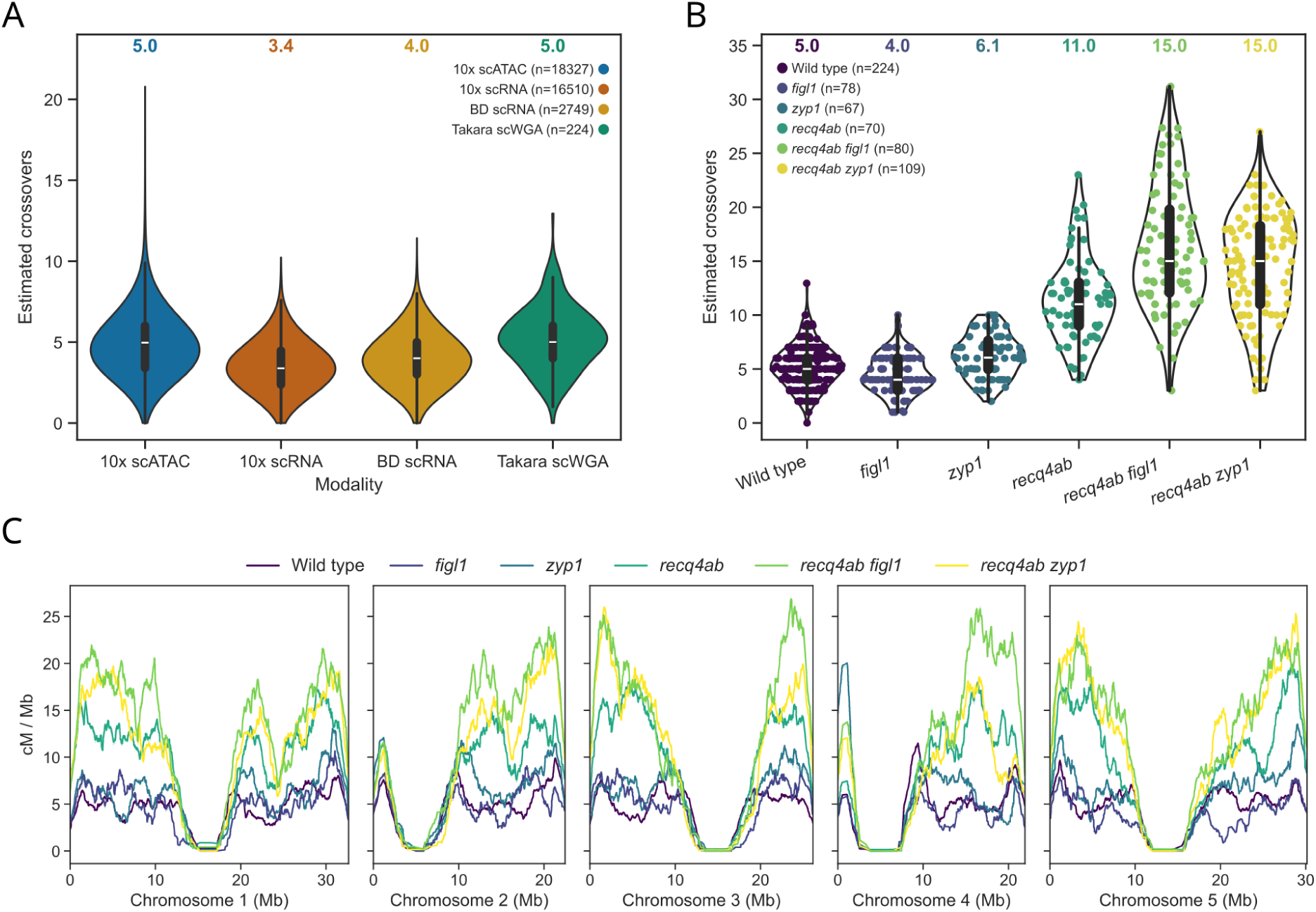
Recombination detection across single-cell sequencing modalities and mutant genotypes. **(a)** Crossover numbers estimated in individual pollen nuclei from each modality. Median crossover numbers per modality are listed above the violin plots. **(b)** Violin plot with overlaid stripcharts, showing the crossover numbers estimated in individual pollen nuclei from different genotypes using the Takara scWGA platform. Median crossover numbers per genotype are listed above the violin plots. **(c)** Recombination landscape of the different genotypes analysed using the Takara scWGA platform, illustrating elevated crossover numbers in mutants throughout the genome.

### Simulation-based evaluation of modality-specific coverage and distribution effects

To quantify how sequencing depth influences crossover detection, we simulated realistic single-cell datasets by projecting known crossovers from progeny-sequencing experiments of wild-type Col-0^♀^ × (Col-0 × L*er*)^♂^ backcrosses (9–11,22,23) onto the empirical fragment distributions of each single-cell modality, adding background noise estimated from the data. Simulated crossover predictions showed a strong dependence on coverage: for example, barcodes simulated from 10x RNA data with fewer than 500 informative fragments had a true positive rate of 71.7%, while those exceeding 500 fragments had a true positive rate of 80.2% (Figure 3A). Telomere-proximal crossovers were disproportionately missed, presumably because the distances between the crossover and the chromosome end are often too short for reliable haplotype inference (Figure 3B). False positive rates were overall low, demonstrating the robustness of the rHMM method to background noise (Figure 3B). For droplet based datasets with lower coverage, the distance between the simulated positions of crossovers and the positions estimated by the model was also greater, illustrating that coverage also affects resolution of crossover localisation (Figure 3C). Consequently, simulated datasets resembling Takara scWGA achieved far higher detection accuracy than droplet-based data, illustrating the trade-off between throughput and sensitivity in single-cell recombination mapping.

**Figure 3.**
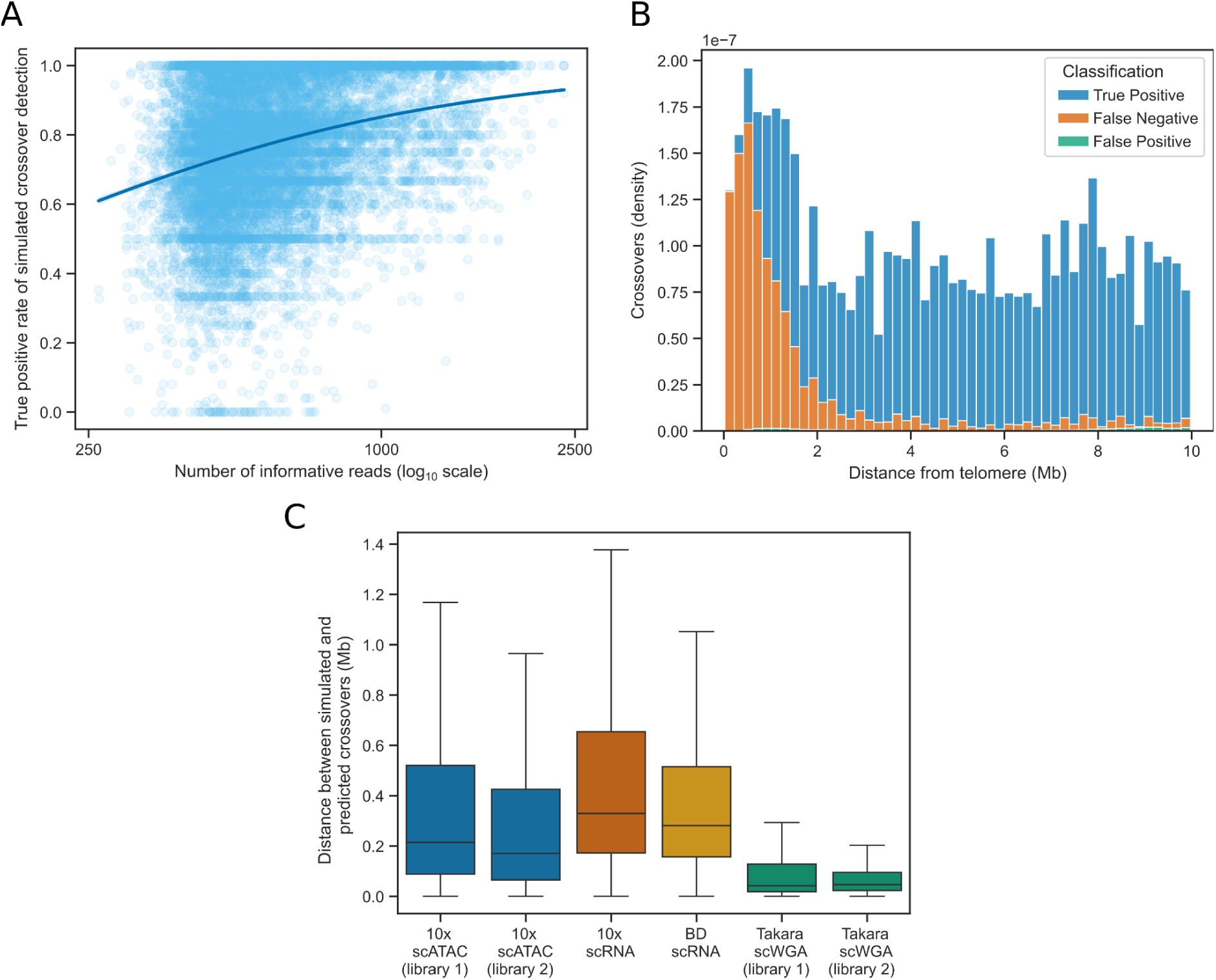
Simulation-based evaluation of crossover statistics. **(a)** True positive rate of crossover detection in barcodes simulated from 10x RNA data, compared to the number of informative reads. **(b)** True positive, false positive and false negative classification of predicted crossovers simulated from 10x RNA data, distributed by distance from the telomere (excluding the beginning of Chromosomes 2 and 4 which contain the Nucleolar Organising Regions). Most undetected (false negative) crossovers occur within 2 Mb of the telomere. **(c)** Boxplot showing the resolution of crossover detection for the different datasets. Plate-based Takara DNA sequencing, which has higher coverage per-barcode, resolves crossover locations more accurately than the lower coverage droplet methods.

### Natural variation in recombination rate and distribution

To test the scalability of coelsch for complex mixtures of genotypes, we profiled pollen nuclei from 40 F₁ hybrids (derived from crossing 20 natural Arabidopsis accessions to both Col-0 and Cvi-0) pooled into a single library using the 10x scRNA modality, which can sample a large number of meioses. Using variant-aware alignment to all 22 parental haplotypes and genotype assignment, 11,534 high-quality nuclei were identified. These nuclei were confidently assigned to 36 F₁ combinations (median = 287 nuclei per hybrid), from which 17 accessions (34 F_1_s) had data for both Col-0 and Cvi-0 crosses. The data from these 34 F_1_s were taken forward for further analysis.

Crossover rate estimates varied widely among accessions, ranging from a median of 3.7 COs / nucleus in Col-0 × Tsu-0 to 7.2 in Cvi-0 × Zin-9 (Figure 4A). Rates were correlated between Col-0 and Cvi-0 crosses with the same accession (Pearson r = 0.71, p = 1.4 × 10^−3^). In most cases, Cvi-0 hybrids exhibited higher overall recombination than the corresponding Col-0 hybrids (Figure 4A). However, the opposite pattern was observed within ∼2 Mb of chromosome termini, with Col-0 hybrids having a higher recombination rate in this region (Figure 4B). To quantify this, we defined a crossover terminality index - the weighted mean number of crossovers where weighting decays exponentially with distance of the crossover from the closest telomere (Figure 4C). This index revealed consistently lower terminal enrichment of crossovers in Cvi-0 hybrids (Figure 4D) compared to their corresponding Col-0 hybrids, implying genetic-background-dependent modulation of crossover localisation.

**Figure 4.**
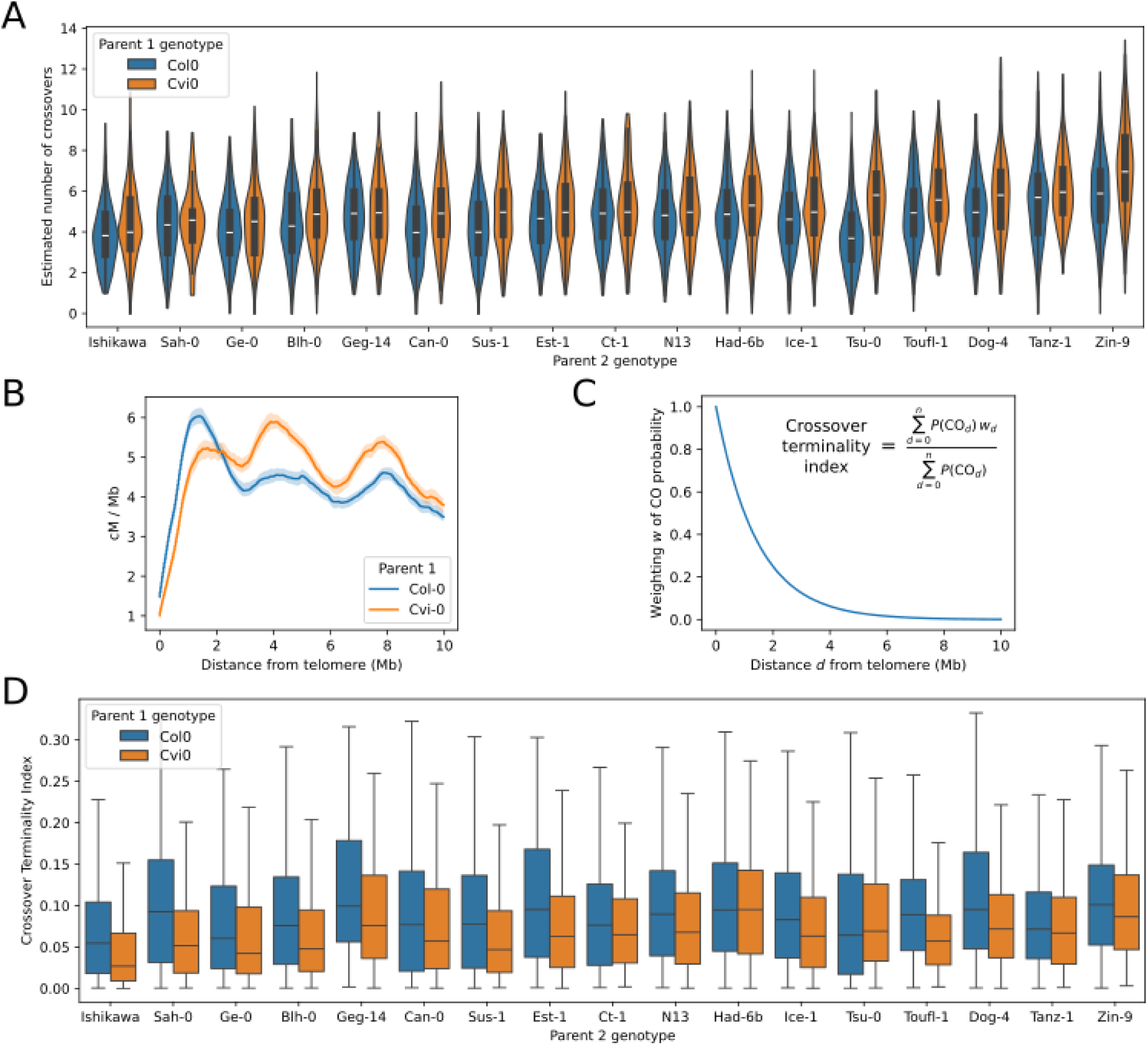
Natural variation in recombination rate and distribution. **(a)** Violin plot showing the estimated number of crossovers per male meiosis for 34 F_1_ hybrids. Violins are trimmed to the observed data range. **(b)** Recombination distribution in cM/Mb at chromosome ends (except the left ends of Chr2 and Chr4 where NORs are present) within 10 Mb of the telomere. Gametes from crosses involving Col-0 have higher recombination rates in telomere-proximal regions than those from crosses involving Cvi-0. Shaded regions show 95% confidence intervals. **(c)** Definition of crossover terminality index, the weighted mean number of crossovers for each gamete, with weightings derived from the distance to the nearest telomere. An exponential decay function with a half life of 1Mb was used. *P(*CO*_d_)* indicates the probability of a crossover at genomic position *d,* whilst *w_d_* indicates the weighting at genomic position *d*. **(d)** Boxplot showing the crossover terminality index for 34 F_1_ hybrids. Despite generally higher overall rates of recombination, crosses involving Cvi-0 tended to have reduced terminal crossovers compared to those involving Col-0.

### Discovery of a large pericentric inversion in Zin-9

While comparing recombination landscapes, we observed a pronounced crossover cold-spot on the right arm of chromosome 2 in gametes from hybrids of the Moroccan accession Zin-9 (24,25), immediately adjacent to the centromere (Figure 5A). To investigate potential structural or epigenomic causes, we generated a *de novo* assembly of the Zin-9 genome using Oxford Nanopore native DNA sequencing. This assembly revealed a previously unrecognised ∼10 Mb pericentric inversion (Chr2:4.5-14.5 Mb) which encompasses the region identified as a recombination coldspot, explaining the absence of crossovers in this interval (Figure 5B). The inversion boundaries coincide with assembly gaps in an earlier Zin-9 genome assembly, indicating that the prior reference-guided assembly of this accession was mis-scaffolded (26).

**Figure 5.**
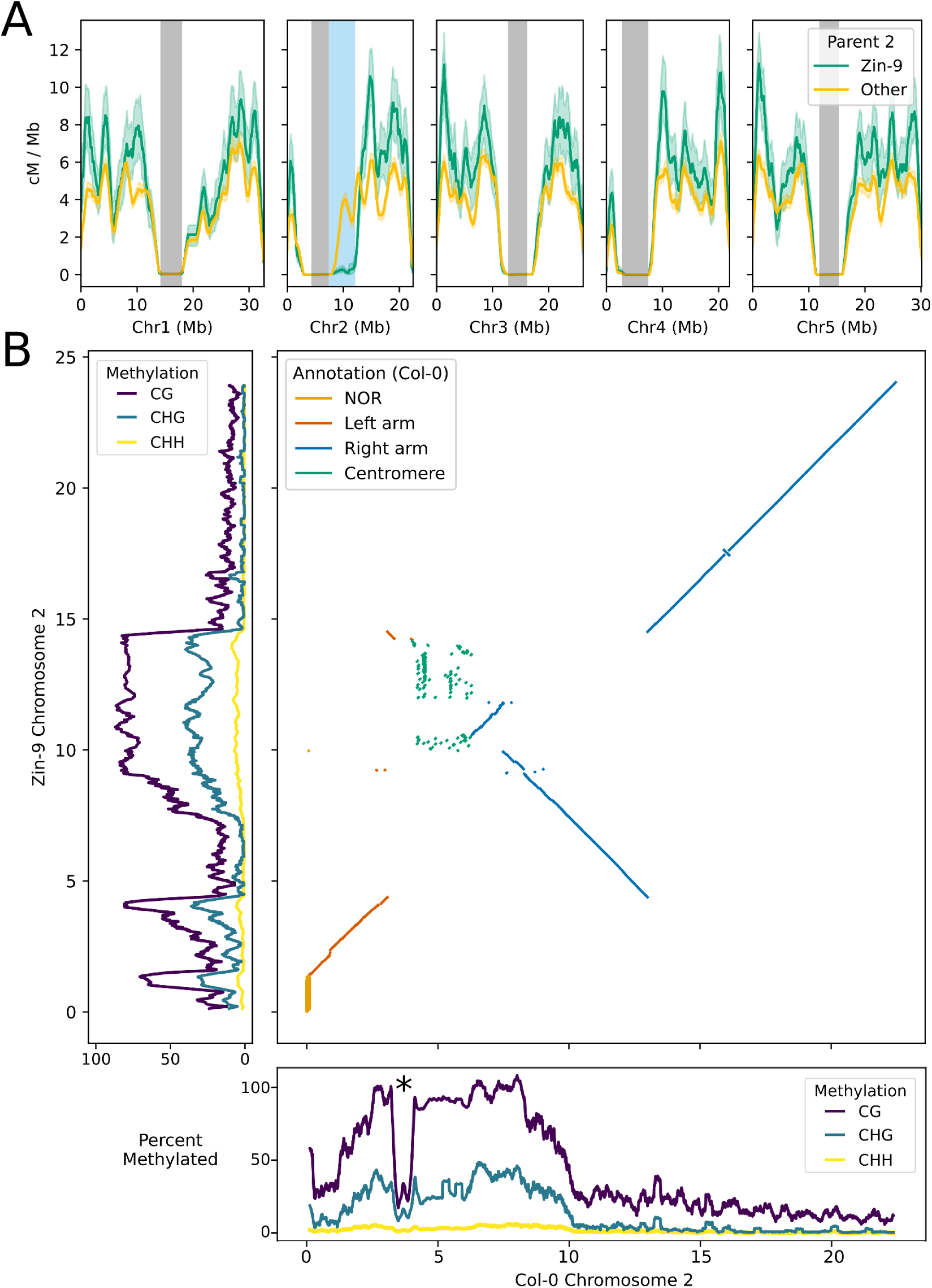
Discovery of a large pericentric inversion in Zin-9. **(a)** Recombination landscapes from 10x scRNA data of pollen from 36 different F_1_ hybrids, comparing hybrids involving Zin-9 to all others. Centromeres are shown as grey shaded boxes. A crossover coldspot specific to Zin-9 F_1_ hybrids on Chromosome 2 (approximate TAIR12 v1 coordinates 8-12 Mb) is highlighted with a light blue shaded box. Shaded regions show 95% confidence intervals. **(b)** Dotplot showing the alignment of a new Zin-9 Chromosome 2 assembly generated using Oxford Nanopore sequencing to the Col-0 CC v1 Chromosome 2 assembly. A 10 Mb pericentric inversion at 4.5-14.5 Mb is clearly visible. Marginal plots show the methylation levels of Col-0 and Zin-9 across Chr2. An asterisk on Col-0 Chromosome 2 marks the position of the mitochondrial DNA insertion, methylation estimates may not be accurate in this region due to unmethylated reads that derive from the mitochondrial genome mapping to the nuclear genome.

5mC methylation profiles showed that the region of the Zin-9 Chromosome 2 at approximately 2.5-4.5 Mb, which is syntenic to the left pericentromere of Col-0, remains heavily methylated and likely heterochromatic, despite being positioned more than 5.5 Mb from the centromere (Figure 5B). Conversely, the centromere–euchromatin junction created by the inversion breakpoint at 14.5Mb on the right side of Zin-9 centromere showed no evidence of heterochromatin spreading (Figure 5B), indicating that chromatin identity is maintained locally rather than determined by linear proximity to centromeric repeats (27–30). Together, these data demonstrate that single-gamete recombination maps can reveal major structural rearrangements and their epigenomic consequences.

## Discussion

Single-cell sequencing of gametes provides a powerful approach for mapping meiotic recombination. Here, we present a unified computational framework that generalises single-gamete haplotyping across droplet- and plate-based modalities, enabling systematic evaluation of their characteristics. By applying coelsch and coelsch_mapping_pipeline to datasets from four different sequencing technologies (13), we recover the expected trends in recombination phenotypes of mutant genotypes (31), resolve crossover landscapes across a number of natural accessions, and discover previously overlooked structural variation (26). This allows informed experimental and analytical choices when designing gamete-based recombination studies, by making the trade-offs between throughput, per-cell resolution, and artefact burden.

Our comparative analyses highlight the trade-offs between droplet- and plate-based sequencing modalities for single-gamete studies. Although only one or two datasets were analysed per modality, and per-dataset summary statistics should therefore be interpreted cautiously, a number of generalisable conclusions can be made. The droplet-based platforms provided an order of magnitude more nuclei per library but with sparser genomic coverage per barcode, while plate-based scWGA yielded fewer nuclei with much higher sequencing depth. The resulting data types capture complementary aspects of the recombination landscape: droplet-based methods are suited to estimating genome-wide crossover frequency distributions across thousands of meioses, whereas scWGA enables higher resolution and accuracy analysis per-individual, in smaller numbers of gametes. The doublet detection framework implemented in coelsch proved essential for both droplet- and plate-base single cell data, confirming that doublet artefacts are a pervasive issue in multiple different modalities (32). Correcting doublet artefacts is necessary to obtain accurate haplotype calls and avoid artificial inflation of crossover counts. These observations show that computational curation, such as that implemented in coelsch, is as critical as experimental optimisation in single-cell recombination mapping.

Simulations using data-derived informative fragment distributions also helped us to determine true and false positive rates of crossover calling for non-doublet barcodes. Although false positives were low, false negatives were more common, particularly near telomeres where short distances between crossovers and chromosome ends limit the information available for detection in sparse data. Although sensitivity is lower than what is achievable with progeny sequencing, relative differences in crossover traits can still be interpreted within a dataset under the assumption that the true-positive rate is comparable across samples.

Beyond technical benchmarking, our results demonstrate the biological applications of single-cell crossover detection. In *A. thaliana* mutants with known perturbations in meiotic recombination, such as *recq4ab*, *zyp1*, and *figl1* (11), our method quantitatively recovered the expected increases or decreases in crossover frequency. The correspondence of crossover landscapes with those determined from previous backcross analyses validates the ability of single-gamete sequencing to measure recombination distributions genome-wide (10,31).

By enabling the simultaneous analysis of dozens of F_1_ hybrid genotypes within a single experimental dataset, our method allows comparative recombination analyses at a scale that has not previously been practical. Applied to natural accessions, this approach identifies substantial variation in crossover rate, consistent with a polygenic quantitative trait (2): rates were correlated between crosses of the same founder genotype with Col-0 and Cvi-0, yet imperfectly so, implying *trans*-acting genetic modifiers and possible non-additive epistatic interactions between haplotypes. The observed variation in crossover terminality further indicates that regulatory differences can shift crossover placement along chromosome arms (4,33).

The consistent absence of crossovers in a defined genomic interval on chromosome 2 also led us to identify a ∼10 Mb pericentric inversion that had been misassembled in the previous Col-0-scaffolded Zin-9 assembly (26). This is the largest natural inversion identified in Arabidopsis to date and suggests that the extent of large-scale structural variation in the species may have been underestimated (26). Such findings highlight the utility of genetic maps in identifying rearrangements that have escaped genome assembly methods. Methylation profiling of the Zin-9 genome further indicated that inversion breakpoints can decouple chromatin state from sequence position, maintaining heterochromatic identity in displaced pericentromeric regions, and an absence of spreading of heterochromatic marks across inversion breakpoints into adjacent euchromatin. These findings are in agreement with previous observations that epigenetic patterns are resilient to major structural rearrangements (27,28,30).

Despite the advances of single-cell methods for crossover analysis, several limitations remain. Coverage heterogeneity remains an intrinsic feature of both scATAC and scRNA data, imposing uncertainty on crossover placement in low-information regions. While doublet detection is effective at removing problematic barcodes, it does so at the cost of wasted sequencing effort, highlighting the need for experimental optimisation to reduce doublet rates. Furthermore, our current approach relies on controlled crossing of isogenic founders with high-quality parental genome assemblies and phased variants; its performance in heterozygous and outcrossing species lacking these resources will depend on reference genome reconstruction and haplotype phasing. Future versions of the coelsch framework could integrate the phasing of variants using genetic linkage information inherent to the gamete-sequencing data to overcome this issue (18,34).

## Conclusions

We present a computational framework for mapping meiotic recombination from diverse single-cell sequencing technologies. By accommodating both droplet- and plate-based approaches, this framework supports analyses that balance the number of gametes sampled with sequencing depth per cell, allowing methods to be matched to specific experimental goals. Our findings underscore the utility of single-cell gamete sequencing for studying the genetic and structural determinants of meiotic recombination (11), uncovering hidden structural variation, and genotyping large populations of recombinants for genetic mapping experiments (12). Meiotic crossover rate is a highly polygenic phenotype exhibiting significant natural variation (2), yet the genetic determinants of much of this variation are still to be discovered. Overall, whilst progeny sequencing may remain the “gold-standard” for high-resolution crossover mapping, single-cell gamete sequencing provides a scalable solution for high-throughput studies of variation in recombination rate and distribution across genotypes, environments, and species.

## Methods

### Plant material and growth conditions

Plant material for benchmarking datasets was generated by crossing Col-0 to L*er*, using Col-0 as the mother. To generate functionally-homozygous single *figl1, recq4ab*, *zyp1,* and double *recq4ab figl1* and *recq4ab zyp1* mutants in a heterozygous Col-0 × L*er* background, crosses were made using *zyp1-1* (8.7.2V1T3) in Col-0 and *zyp1-6* (1.12V5T2) in L*er* (22), *recq4a-4* (GABI_203C07) in Col-0 (35), *recq4a-W387X* in L*er* (36), *recq4b-2* in Col-0, *figl1-19* (SALK_089653) in Col-0 (11) and *figl1-12* in L*er* (37). For the diversity panel of 40 different accessions, crosses were made between either Col-0 or Cvi-0 and 20 different natural accessions (Blh-1, Can-0, Ct-1, Dog-4, Est-1, Ge-0, Geg-14, Had-6b, Ice-1, Ishikawa, Jea, Kz-9, N13, Pyl-1, Sah-0, Sus-1, Tanz-1, Toufl-1, Tsu-0, Zin-9) acquired either from the Versailles Collection (http://publiclines.versailles.inra.fr/) or from Angela Hancock (MPIPZ). These 20 accessions were selected from a group with chromosome-scale genome assemblies (26), to represent a cross section of global genetic diversity and maximise variants that could be used for genotyping/demultiplexing of single cell barcodes.

Plants were grown in growth chambers with a photoperiod of 16 h of light and 8 h of darkness and 22°C daytime and 18°C night time temperatures. After flowering, F_1_ pollen was collected from mature anthers just prior to anthesis, flash-frozen in liquid Nitrogen and stored at –80°C.

### Nuclei isolation for 10x scATAC

For 10x scATAC-seq, pollen was isolated from approximately 600 flowers (Library 1) or 900 flowers (Library 2). Frozen flowers were submerged in LB01 buffer, modified from Doležel et al. (38), consisting of 15 mM Tris-HCl pH 7.5, 2 mM EDTA pH 8, 0.5 mM spermine·4HCl, 80 mM KCl, 20 mM NaCl, 5 mM 2-mercaptoethanol, and 0.15% Triton X-100, and vortexed for 60 seconds. The suspension was filtered through a 100 µm Celltrics filter placed above a 5 µm filter for pollen collection. Pollen retained on the 5 µm filter was mechanically disrupted using a plastic pestle to release nuclei, which were washed with 350 µl LB01 buffer. This filtering and grinding procedure was repeated until all pollen was processed.

Nuclei were isolated using Nuclei Isolation Buffer (NIB), modified from Sikorskaite et al. (39), consisting of 10 mM MES-KOH pH 5.4, 10 mM NaCl, 10 mM KCl, 2.5 mM EDTA pH 8, 250 mM sucrose, 0.1 mM spermine·4HCl, 0.1 mM spermidine, and 1 mM DTT, with 1% BSA Fraction V added immediately before use. NIB was used to prepare a 35% v/v Percoll solution.

Density gradients were prepared in 15 ml tubes by layering 2.5 M sucrose beneath 35% Percoll. For Library 1, 750 µl sucrose and 3 ml Percoll were used, and for Library 2, 2 ml sucrose and 6 ml Percoll were used. Isolated nuclei were gently layered onto the Percoll–sucrose gradients and centrifuged for 35 min at 3,500 g and 4 °C with acceleration 4 and deceleration 2. Nuclei were collected from the Percoll–sucrose interface, filtered through a 5 µm filter, and washed twice by centrifugation for 6 min at 1,000 g and 4 °C.

Final nuclei pellets were resuspended in 10 µl 1× nuclei buffer from 10x Genomics supplemented with 1% BSA by gentle pipetting. Nuclei quality was assessed by DAPI staining and fluorescence microscopy, and concentration was estimated by propidium iodide staining using a Luna-FX cell counter.

### Nuclei isolation for Takara scWGA

For Takara scWGA, pollen from 70–200 flowers per genotype (including meiotic mutants) was used. Pollen nuclei were collected from each genotype independently in NIB as described for 10x scATAC, omitting Percoll purification. Nuclei quality and concentration was manually estimated by propidium iodide staining under a microscope.

### Nuclei isolation for 10x and BD scRNA

For scRNA-seq using 10x Genomics and BD platforms, approximately 400 flowers per hybrid genotype were pooled into a 50 ml tube containing 10 ml pre-chilled scWPB, modified from Loureiro et al. (40), consisting of 0.2 M Tris-HCl, 3 mM MgCl₂·6H₂O, 0.1 mM EDTA·Na₂·2H₂O, 86 mM NaCl, 10 mM Na₂S₂O₅, 1% PVP-10, 0.5 mM spermine·4HCl, and 0.5 mM spermidine. Pollen was released by vortexing and filtered through a 100 µm filter placed on top of a 10 µm filter. Pollen retained on the 10 µm filter was ground with a plastic pestle to release nuclei, which were washed in scWPB supplemented with 5 mM DTT, 2% BSA Fraction V, and 0.2 U/µl Protector RNase Inhibitor.

Extracted nuclei were stained with 1 µg/ml DAPI and sorted using a BD FACSAria Fusion with a 70 µm nozzle and 0-16-0 precision. Approximately 200,000 nuclei were collected per library. Sorted nuclei were collected into 1× PBS containing 1% BSA and 0.2 U/µl Protector RNase Inhibitor, with collection volume scaled at 1 µl per 1,000 sorted events. Nuclei concentration and quality were assessed prior to library preparation.

### Single-cell library preparation

Single-cell libraries were prepared using commercial kits according to the manufacturers’ instructions. 10x Genomics scATAC libraries were generated using the Chromium Next GEM Single Cell ATAC v2 kit. 10x Genomics scRNA libraries were prepared using the Chromium GEM-X Single Cell 3′ Reagent Kit v4. BD scRNA libraries were prepared using the BD Rhapsody cDNA and WTA Amplification kits. scWGA libraries were generated using the Takara Shasta Whole Genome Amplification Kit for the ICELL8 cx system. All single-cell libraries were sequenced by BGI using DNBSEQ G-400 technology.

### Variant-aware mapping

Reads were aligned using coelsch_mapping_pipeline (v0.1.1), a Snakemake-based workflow for haplotype-aware alignment of single-cell datasets. For each dataset, the assemblies of the founder haplotypes were first aligned to the Col-0 CC v1 (GCA_028009825.1) reference genome using minimap2 with preset asm10 and z drop 500 (v 2.30) (41). For Col-0 × L*er* datasets, only the L*er* assembly (GCA_946406525.1 (42)) was aligned to Col-0; for the diversity panel, assemblies of 21 non-reference founders (Cvi-0 + 20 diverse accessions) were aligned (26).

Core-syntenic regions were identified using syri (v 1.7.1) (43) and msyd (v 0.30) (44). Within these core-syntenic regions, SNPs and short indels (< 50 bp) were extracted as markers for variant-aware mapping. Low-mappability and repetitive regions were identified in each assembly using genmap (v 1.3.0) (45) with a kmer size of 100 and an edit distance of two. Any variants contained within non-unique regions under these parameters were excluded from marker sets to avoid ambiguous alignment (46). For Col-0 × L*er*, the retained core-syntenic regions covered 108.6 Mb of the Col-0 TAIR12 v1 genome with 742,839 marker variants; for the diversity panel, core-syntenic regions represented 86.5 Mb of the Col-0 CC genome, with an average of 612,522 marker variants per haplotype, relative to Col-0 (range [491,717-812,880]).

Per-haplotype variant sets were then applied to the Col-0 CC reference sequence using STAR (version 2.7.11a) in consensus-indexing mode to generate haplotype-specific genome indices (19,20). Reads were aligned to each haplotype-transformed index using STAR or STARsolo (47), depending on library type. For droplet-based single-cell libraries (10x Genomics, BD), alignments were performed with STARsolo using the corresponding barcode whitelists; for plate-based libraries, reads for each barcode were aligned independently with STAR. Spliced alignment was enabled for scRNA-seq datasets (maximum intron = 20 kb) and disabled for scATAC-seq and scWGA datasets. For paired ended data, a max alignment gap between mates of 500bp was used.

Following alignment, BAM files were converted back into Col-0 CC coordinate space using STAR consensus mapping to maintain a common reference frame (20). For each read, the haplotype (or multiple haplotypes) producing the highest alignment-score (AS tag) was identified, and these haplotype-specific alignments were retained to generate per-barcode marker profiles for downstream recombination and genotype analysis. We refer to sequenced fragments overlapping one or more parental marker variants as informative fragments.

### Data loading and genotyping of parental accessions

Per-barcode haplotyping data was loaded into a custom json format using coelsch (v 0.6.1). After assigning barcode-level marker counts to parental haplotypes, we summarised alignments into 25 kb bins across the genome. For the diversity panel containing 40 possible F_1_ hybrid genotypes, each barcode was genotyped using an expectation-maximisation (EM) algorithm under a simple error model (12). Each genotype was represented as a pair of parental haplotypes, and each observed marker was treated as supporting either one of these haplotypes (a true match) or neither (a mismatch, explained by an estimated global error rate).

The EM algorithm iteratively updated posterior probabilities for all 40 candidate genotypes and the global error rate until convergence, defined as a total change in probability < 0.001 or a maximum of 1000 iterations. To assess assignment uncertainty, 25 bootstrap resamplings were performed per barcode by drawing markers with replacement. Genotype probabilities were recomputed for each bootstrap replicate, and the mean posterior probability was taken as the final genotype confidence score. The genotype with the highest mean probability across bootstraps was assigned to that barcode.

### Data cleaning

After assigning each barcode to a parental genotype, informative alignments distinguishing the two parental haplotypes were identified and used to construct marker profiles for recombination analysis. Marker profiles were normalised using a per-chromosome, quantile-based shrinkage procedure that dampens extreme coverage values arising from highly expressed genes (in scRNA data) or from regions of high marker density and collapsed repeats (in the general case). To mitigate noise from alignment errors, ambient nucleic acids, and non-syntenic markers, two filtering procedures were applied:

Haplotype imbalance filter: For each parental genotype, haplotype-specific read distributions were aggregated across all barcodes to detect bins exhibiting systematic allelic imbalance. For each bin, the proportion of reads assigned to each haplotype was compared to the expected 50:50 ratio. Bins showing extreme distortion, where one haplotype accounted for more than 75 % of reads among bins supported by at least 20 barcodes, were classified as artefactual and masked. This filter removes regions affected by mapping bias, local copy-number variation, or collapsed repeats that distort haplotype ratios.

Background subtraction filter: For each barcode, background contamination was estimated using a convolution-based masking approach that identifies the locally dominant haplotype (foreground) and treats counts for the alternative haplotype as background noise. The per-barcode background fraction was computed as the proportion of total counts assigned to the background after smoothing over a fixed genomic window. For each genotype, a normalised background profile was then constructed by averaging background counts across barcodes. Barcode-specific background signal was subsequently subtracted deterministically from marker counts in proportion to both the estimated background fraction and the genotype-level background probability model, ensuring non-negative corrected values. This procedure removes systematic sources of noise such as ambient RNA and recurrent mapping artefacts.

### Crossover detection

To infer haplotype state probabilities along individual barcodes, we implemented a rigid Hidden Markov Model (rHMM) that constrains recombination transitions to occur only at fixed genomic intervals, approximating crossover interference at adjacent loci. Modelling was performed using pomegranate (v 1.0.0) (48) and PyTorch (v 2.1.2) (49). Three model configurations, for haploid (two-state), diploid backcross (two-state), and diploid F₂ (three-state) data are currently available. In all cases, states correspond to haplotypes, or haplotype combinations, and emission parameters can be either estimated from the data or supplied manually. Each haplotype combination is represented as a fixed linear chain of length r, corresponding to the minimum number of genomic bins between allowable transitions (8,12). Transition probabilities are rigidly structured such that each position in the chain can either self-loop, progress deterministically to the next position, or, at chain boundaries, transition to a different haplotype state representing a crossover event.

Emission probabilities at each chain position of the rigid Hidden Markov Model (rHMM) were estimated directly from the marker count data. For haploid datasets, foreground and background Poisson means, along with the zero-inflation fraction, were inferred using a two-component mixture model consisting of a Dirac delta (empty bins) and Poisson emissions. For diploid datasets (backcross individuals), parameters were estimated using an extended mixture including heterozygous and homozygous Poisson components, with foreground and background haplotypes identified via local smoothing and column re-ordering. Transition parameters were defined globally from the average expected recombination rate of Arabidopsis (∼4.5 cM/Mb) (10) and genomic bin size (25 kb). The rigid chain length and terminal length were set as 1 Mb and 50 kb, respectively.

For each barcode, haplotype probabilities were inferred by applying the trained rHMM to the per-bin informative fragment count matrices. Predictions were run chromosome by chromosome in batches of up to 128 barcodes with parallel threading. The model outputs per-bin posterior probabilities for each haplotype state, which were collapsed into single-haplotype probabilities representing the local likelihood of the alternative parental haplotype. These probability profiles were stored as json objects for subsequent recombination analysis and visualisation.

### Doublet prediction

Doublet barcodes were identified using a simulation-based classification approach (32). Synthetic doublets were generated by randomly combining pairs of real barcodes of similar coverage. These simulated doublets were processed identically to real barcodes: haplotype probabilities were predicted using the trained rHMM, and the agreement of the simulated informative fragment distributions and predicted haplotype probabilities were calculated as a measure of data quality. A two-component Gaussian mixture model was fitted to these metrics to separate doublet and singlet barcodes, using parameters from the simulated data as the starting point for the doublet distribution. Predicted probabilities were used to flag putative doublets for downstream filtering.

### Simulation of recombination events

To evaluate model performance, synthetic single-cell marker datasets were generated using a simulation framework, also implemented in coelsch. Ground-truth haplotypes can be provided either as BED-formatted interval files or using coelsch’s JSON-formatted prediction records. For each chromosome, a per-bin background signal and barcode-specific contamination rate were estimated from the input dataset using the same convolution-based procedure applied during background correction. Simulated singlet barcodes were then created by applying the known ground-truth haplotype (10,11) to real informative fragment profiles. Foreground and background signals were randomly sampled from these profiles using the estimated background model as weights, and counts were reassigned to the appropriate haplotype according to the ground truth allele state for each bin. This procedure creates data with known crossover patterning while preserving the empirical noise structure and per-barcode coverage distribution of the original data. Optionally, doublet barcodes can also be simulated by randomly pairing and summing marker profiles from two real barcodes of similar coverage, using the same method as described in the doublet prediction section.

### Analysis of recombination rate and distribution

Crossover frequency was quantified for each barcode as the summed absolute difference in haplotype probabilities between adjacent genomic bins, after applying a minimum per-bin transition probability threshold of 5×10⁻³ to reduce the impact of noise. Note that this method can produce “fractional” crossovers when the model is uncertain. To estimate recombination landscapes, haplotype probability gradients were calculated across all barcodes and smoothed with a rolling mean over 1 Mb windows. Recombination rate was expressed in centimorgans per megabase (cM / Mb) by normalising the summed crossover probabilities to the effective number of barcodes per position and scaling by genomic distance. To account for lack of marker density near chromosome termini, effective barcode counts were derived from the range of non-zero marker coverage. For each genotype, 100 bootstrap resamples of barcodes were used to estimate uncertainty in the recombination landscape. All analyses were performed using coelsch’s recombination submodule.

Crossover terminality index was defined as the weighted fraction of crossover probabilities in proximity to the telomeres of each Chromosome. Weightings were generated using an exponential decay function over each haplotype bin, with a half life of 1 Mb.

### Nanopore sequencing and genome assembly of Zin-9

High-molecular-weight genomic DNA from *Arabidopsis thaliana* accession Zin-9 was sequenced using Oxford Nanopore native DNA sequencing on a PromethION P2 solo platform. Raw reads were basecalled with Dorado (v 1.1.1) using the SUP model with 5mC modification calling enabled. De novo assembly was performed with hifiasm (v 0.25.0) (50) using parameters --ont -l0 --rl-cut 10000 --sc-cut 15. Assemblies were scaffolded against the Col-0 CC reference genome using RagTag (v 2.1.0) (51). Whole genome alignment of Zin-9 to Col-0 was performed using minimap2 (v 2.30) with preset asm10 (41). 5mC methylation frequencies were extracted and summarised across the assembly using modkit (v 0.5.0) and bedtools (v 2.30.0) (52).

### Workflow management

All pipelines were written and executed using Snakemake (53), with Python scripts and the Scientific Python stack (54–57), as well as analysis using Jupyter notebooks and the Jupyterlab environment (58). Figures were created using BioRender and Inkscape.

## Data and code availability

Sequencing data and assemblies generated for this study are available from ENA project PRJEB107034. The Col-0 TAIR12 (Col-CC) assembly version 1 is available from NCBI GenBank accession GCA_028009825.1, and the L*er* assembly from GCA_946406525.1. Col-0 × (Col-0 × L*er*) backcross datasets used for simulations are available from ENA projects PRJNA268060, PRJEB79761, PRJEB49306, PRJEB52714, PRJEB40479 and PRJEB49306. The genome assemblies of all other accessions used in the diversity panel are available from the MPI Edmond repository DOI:10.17617/3.AEOJBL. coelsch and coelsch_mapping_pipeline are available on GitHub at https://www.github.com/schneebergerlab/coelsch and https://www.github.com/schneebergerlab/coelsch_mapping_pipeline respectively.

## Author contributions

KS and RM conceptualised the project. KS, RM and MP supervised the project. SA, BW, VS, MM and BH generated the data. MP, JF and XD designed and implemented the bioinformatic pipelines and analyses. MP and JF wrote the manuscript with input from all authors.

## Acknowledgements

We are grateful to the NGS Facility for Integrate Genomics at the Universitätsmedizin Göttingen for the use of their Takara iCELL8 machine, and to Samia Oussous from Takara Biosciences for her support in generating the Takara scWGA libraries. We also thank Craig Dent for helpful comments on the manuscript.

## Supplementary figures

**Supplementary Figure 1.**
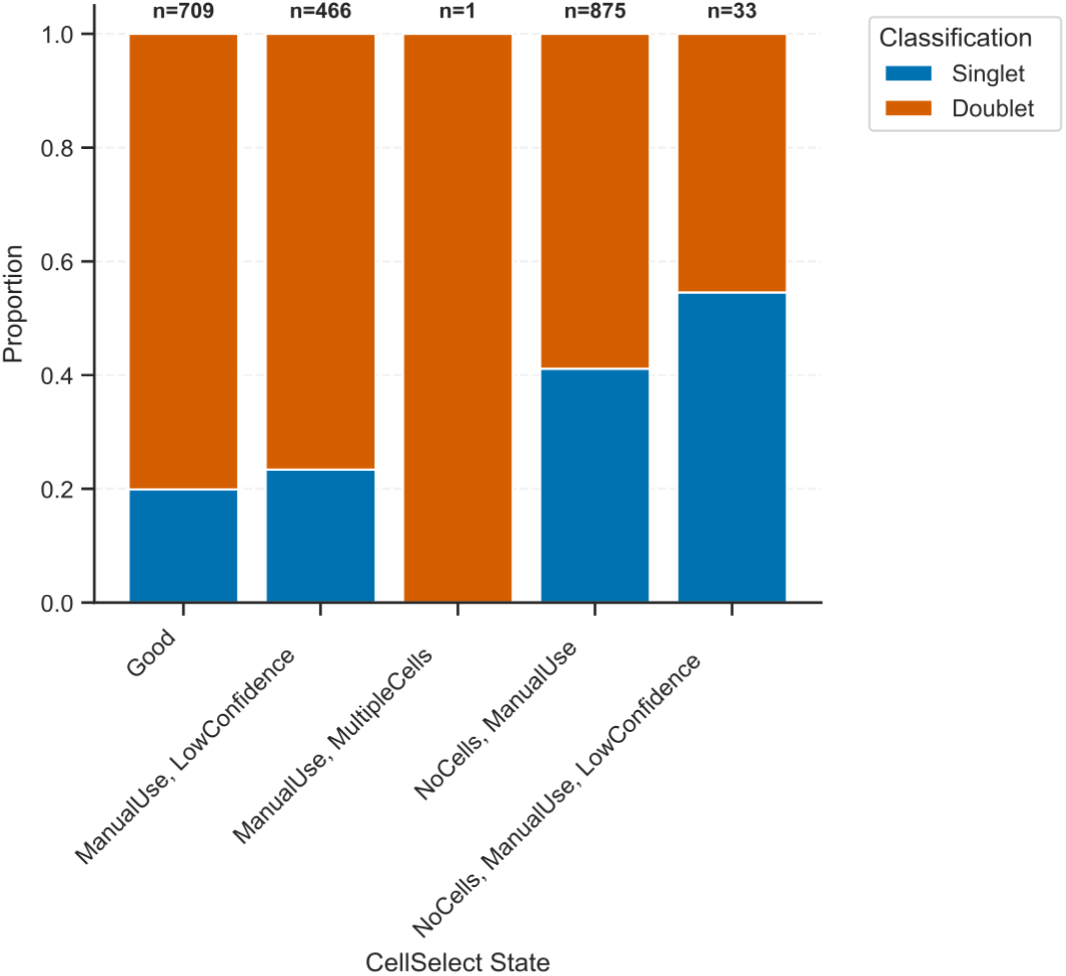
Proportion of singlet and doublet barcodes across CellSelect states. Barplot showing the proportion of singlet and doublet barcodes per category assigned by CellSelect. CellSelect states were assigned by the CellSelect software to identify single-nucleus wells in Takara’s iCELL8 single-cell system. Doublets were defined by filtering barcodes with a doublet probability score < 0.5 using coelsch.

